# Longitudinal Saliva Omics Responses to Immune Perturbation: A Case Study

**DOI:** 10.1101/2020.06.16.156133

**Authors:** George I. Mias, Vikas Vikram Singh, Lavida R.K. Rogers, Shuyue Xue, Minzhang Zheng, Sergii Domanskyi, Masamitsu Kanada, Carlo Piermarocchi, Jin He

**Affiliations:** Michigan State University, Biochemistry and Molecular Biology, East Lansing, MI 48824, USA; Michigan State University, Institute for Quantitative Health Science and Engineering, East Lansing, MI 48824, USA; Michigan State University, Microbiology and Molecular Genetics, East Lansing, MI 48824, USA; Michigan State University, Physics and Astronomy, East Lansing, MI 48824, USA; Michigan State University, Pharmacology and Toxicology, East Lansing, MI 48824, USA

**Author notes:** these authors contributed equally to this work (George I. Mias, Vikas Vikram Singh, Lavida R.K. Rogers).

## Abstract

Saliva omics, a rapidly developing field for non-invasive diagnostics, may be utilized for monitoring very young or elderly populations, as well as individuals in remote locations. In this study, multiple saliva omics from an individual were monitored over 100 timepoints, over three periods involving: (i) hourly sampling over 24 hours without intervention, (ii) hourly sampling over 24 hours including immune system activation using the standard 23-valent pneumococcal polysaccharide vaccine, (iii) daily sampling for 33 days profiling the post-vaccination response. At each timepoint total saliva transcriptome and proteome were profiled, and salivary extracellular vesicles were derived, from which small-RNA sequencing was used to determine RNA, miRNA, piRNA and bacterial RNA components. The two 24-hour periods were used in a paired analysis to reveal vaccination responses. Temporal trends were classified and collective behavior revealed broad immune-responses captured in saliva, both at the innate as well as the adaptive response time frames.

## Introduction

Precision medicine continues its rapid development toward clinical applications aided by new sequencing technology and computational capabilities. Major efforts have concentrated on evaluating disease risk from genomic information^1, 2^, including direct to consumer platforms, like 23andMe^3^, as well as pharmacogenomic evaluations^4^. Implementing omics profiling in the clinic will require evaluation of patients over time, and the utility of such profiling has been evaluated in individual monitoring pioneered in the integrative Personal Omics Profiling (iPOP) study^5^, and expanded recently to include profiling using electronic health devices^6^, the Pioneer study^7^ (which also incorporated behavioral coaching to improve clinical biomarkers based on participants’ individual data), utilizing host-microbiome data in insulin resistant individuals in a study of weight gain^8^ and in prediabetics^9^, investigating biological age^10, 11^ as well as monitoring of astronauts in the recent NASA twin study^12^.

In this investigation we are extending integrative omics to evaluate the utility of such monitoring using saliva. There has been long-standing interest in saliva for non-invasive diagnostics and health monitoring, and saliva omics is an emerging field, with broad profiling that includes total saliva RNA and proteomes, as well as cell-free RNA identification, extracellular vesicle (EV) profiling, miRNAs as biomarkers, and salivary microbiomes^13–25^. Utilizing of saliva for non-invasive monitoring is important in evaluating vulnerable populations, including infants, children, older adults and immunocompromised individuals. Additionally, saliva is important in evaluation of health in remote or underserved locations, when limited resources are available, where processing of blood samples might not be feasible, or a physician may not even be available. Such monitoring is also of particular interest for evaluating active personnel, including astronauts in deep space missions. The recent twin astronaut study evaluated multi-omics utility but also highlighted the logistic issues of using blood samples when these cannot be processed on-site^12^. The COVID-19 pandemic has additionally ignited interest in the use of saliva for rapid diagnostics, towards a rapid and minimally invasive diagnostic that can be used without risk to personnel (possibly as a home-use kit), including profiling viral loads (from posterior oropharyngeal samples)^26^, and current work continues to evaluate the sensitivity of saliva for practical implementations^27–30^.

We are carrying out a clinical trial monitoring individualized response to pneumococcal vaccination, and in a proof-of-principle case-study, we monitored individualized response to the standard 23-valent pneumococcal polysaccharide vaccination (PPSV23), in a generally healthy individual (Caucasian male 38, has reported chronic sinusitis), and carried out integrative profiling on saliva pre- and post- vaccination with pneumococcal PPSV23 vaccine. This is to our knowledge the most extensive saliva-focused omics dataset on an individual, covering 104 timepoints over one year. The period covers a healthy period as well as monitoring of innate and adaptive immune responses following vaccination. Protein and RNA from saliva were produced over 100 timepoints over the course of 1 year, and comprehensive untargeted proteomics and RNA-sequencing were carried out for all samples. The saliva sampled timepoints included three periods of particular reference in this manuscript: (i) 24 hours hourly sampling during to assess a healthy hourly baseline (ii) 24 hours hourly sampling that included vaccination with pneumococcal vaccine (PPSV23) to assess response to the vaccine (iii) daily sampling following the vaccination to assess potential innate and adaptive immune responses reflected in the molecular saliva components.

Our study reveals multiple changes in response to pneumococcal vaccination that are observable in saliva. The microscopic collective behavior of multiple omics reflects physiological changes associated with immune response, including fever, innate and adaptive responses profiled over multiple scales. This case study provides a resource for future saliva studies, towards more effective non-invasive diagnostics.

## Results

### Samples and Assays

We followed a single individual (m, 38, Caucasian), in general good health (has reported chronic sinusitis) over the span of a year. To observe whether the effects of perturbation can be profiled in saliva we carried out the profiling over 3 time frames. In the first 24 hr time frame (TFH1) we established a baseline, obtaining a saliva sample from the subject hourly without perturbation over his standard routine. In the second 24 hr time frame (TFH2), the subject was vaccinated with pneumococcal polysaccharide vaccine (PPSV23) within 3.5 hrs of waking up (at 10.30 am), while otherwise maintaining a similar routine as in the first period (including food intake and meal timing), and again saliva samples were taken hourly. We should note here that the subject reported fever ~7.5 hours post the vaccination (between timepoints at 5 and 6 pm), lasting for about 4 hrs (10 pm). The two time periods, TFH1 and TFH2, were treated as paired and combined in the analysis below (TFΔ) to identify changes induced by the vaccination, by effectively removing daily normal routine effects for this individual. Additionally, in the third time frame (TFD) we monitored the subject daily for over a month, pre- and post- vaccination to identify potential immune changes over both innate and adaptive time frames, Fig. 1.

**Figure 1.**
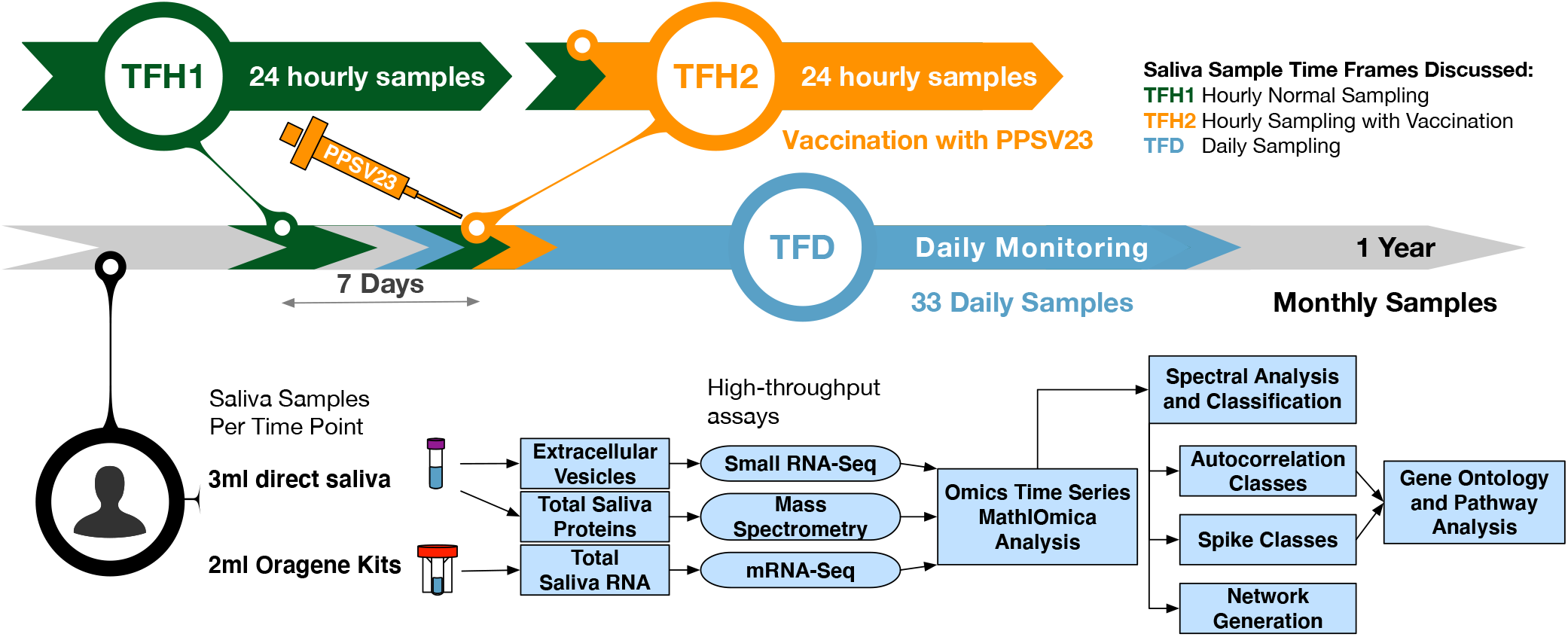
Study Overview. **a** The study followed an individual over the span of a year, collecting 100+ saliva samples. In this manuscript we discuss three time frames of interest for saliva sampling: (i) TFH1 where 24 hourly samples were taken, (ii) TFH2, where 24 hourly samples were taken but the subject was also vaccinated with PPSV23 during this time frame, (iii) TFD where daily samples were taken. **b** At each timepoint saliva were used to extract extracellular vesicle RNA, total saliva RNA and protein for RNA-seq and proteomics. The data was used to generated time series with `MathIOmica`, which revealed multiple trends corresponding to response to immunization with PPSV23.

The daily samples were all taken at 8 am, to limit variability. Saliva was sampled both for downstream total RNA profiling, mass spectrometry proteomics, as well as for extraction of extracellular vesicles which were profiled for a variate of small RNA molecular species (both host and non-host).

### Analysis of Total Saliva and Time Series Constructions

#### Total Saliva Transcriptomics

We profiled the transcriptome from total saliva at all timepoints using RNA-sequencing (RNA-seq) on extracted mRNA (150bp paired-end reads, stranded). We mapped the RNA-seq data using `Kallisto`^31, 32^, and adjusted the values across timepoints using `sleuth`^33^(`DESeq`^34^ adjustment of Transcripts per Million[aTPM], resulting in 81,098 GENCODE^35^ annotation transcripts showing non-zero values for at least 1 timepoint). We carried out downstream analysis in `MathIOmica`^36^, and selected the different timepoints for each of the three time frames. We tagged 0 aTPM values as Missing, filtered for noise (aTPM < 1), removed transcripts with more than 1/4 timepoints missing (i.e reported as zero), and transcripts with constant values across all timepoints, to finally obtain 15,621, 7,493, and 8,155 transcript time series of aTPM values for TFH1, TFH2, and TFD respectively. All values were normalized to a reference - using the first timepoint for TFH1 and TFH2 and the pre-vaccination timepoint for TFD. We then calculated the TFΔ values by calculating the differences per transcript between TFH1 and TFH2 to obtain 7,311 TFΔ time series (after removal of transcripts not overlapping across TFH1 and TFH2).

#### Total Saliva Proteomics

We profiled the total saliva proteome, using isobaric tandem mass tags (TMT) for quantitation using LC-MS/MS (liquid chromatography followed by mass spectrometry). We identified 12,473 proteins overall, with 11,005 proteins (UniProt identifiers^37^) based on 1 unique peptide per protein, (4,141 proteins based on 2 unique peptides per protein) overall across all 95 samples where proteomics was carried out. Relative protein intensities were computed against a common pooled sample comprising of multiple healthy (pre-vaccination) weekly samples that was used across all TMT sample pools. The data were thus combined, and normalized using a Box-Cox^38^ transformation to obtain normal distributions. To construct the time series, the data were filtered again for 1 unique peptide, having less than 1/4 missing values, and no constant time series to obtain 1,227, 1,547, 1,046, and 1,209 proteomics time series for TFH1, TFH2, TFΔ and TFD respectively. All timepoint intensities were defined with respect to the first timepoint intensity for the hourly series for each respective protein, and to the vaccination day for TFD.

#### Analysis of Saliva Extracellular Vesicles

In addition to considering total saliva, we also implemented consistent extraction of EVs from 1ml saliva, using ExoQuick-TC (SBI) and an overnight precipitation to obtain EV pellets, from which we extracted RNA. We carried out nanoparticle tracking analysis (ZetaView, Particle Metrix) and recorded median concentrations of 6.2 × 10^10^ particles/ml with EV peak of 114.5±4 nm, Fig. 2.

**Figure 2.**
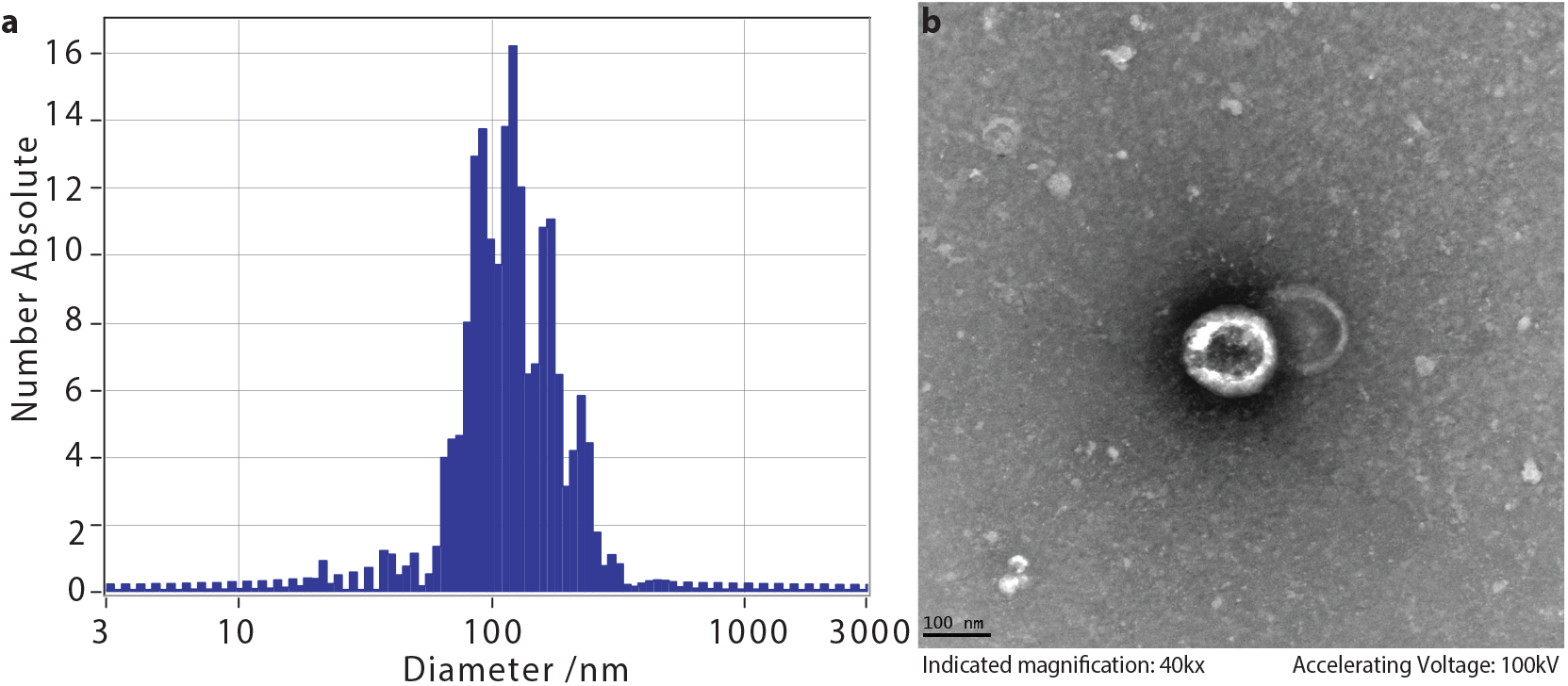
Extracellular vesicle profiles. **a** The EV size was profiled using ZetaView (Particle Metrix) with median concentrations of 6.2 × 10^10^ particles/ml with EV peak of 114.5±4 nm. **b** EVs were imaged using transmission electron microscopy.

The extracted EV RNA was sequenced (small RNA-sequencing) and the results were mapped to multiple databases using the `exceRpt`^39^ through genboree.org^40^. These included GENCODE transcripts, PIWI interacting RNA (piRNA), micro RNA (miRNA), and exogenous genomes and exogenous ribosomal RNA (rRNA) genomes, and multiple other biotypes, Fig. 3. The various biotypes detected have different variabilities, Fig 3a. The majority of detected reads are from exogenous genomes (~10^6^) and protein coding (GENCODE transcripts, 106 as well). The different biotypes per sample, Fig 3b, for the TFH1, TFH2 and TFD time frames are indicative of a change in the relative distributions of the biotypes for the daily samples following the vaccination (increase of exogenous genomes content). This may be partly attributed to sampling as all TFD samples were taken at 8 am, and the corresponding early samples for TFH1 and TFH2 are also similar in biotype relative abundances, but differ in later samples during each day.

**Figure 3.**
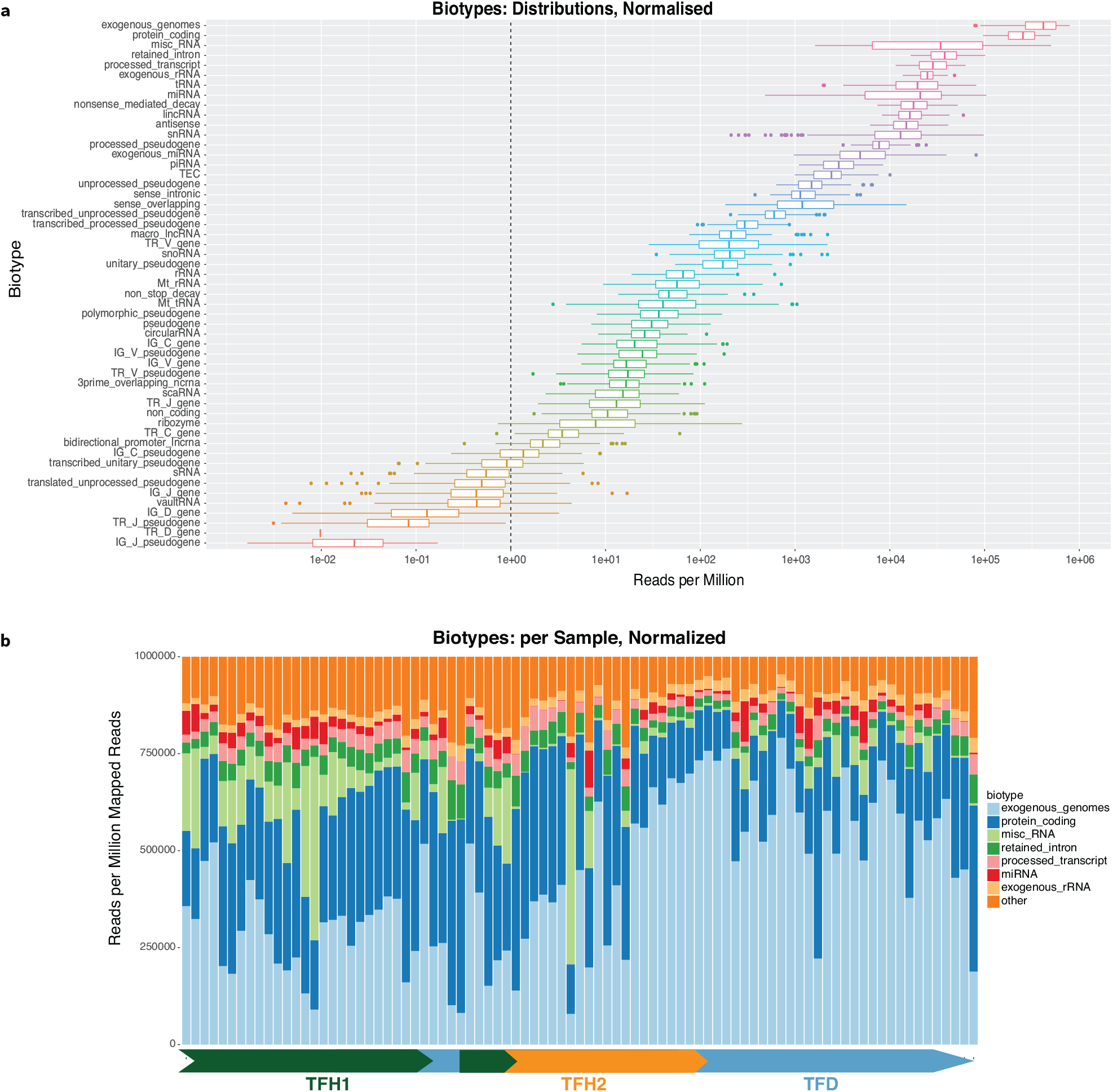
Extracellular vesicle biotypes. **a** Various biotypes were detected in EVs, with the overall distributions as shown (results from `exceRpt`^39^ mapped small-RNA-seq from Saliva (~7 20M reads/sample, 50bp/read).**b** The distributions of hourly and daily samples showed variability, particularly in the reduced exogenous genome content in the hourly samples.

In terms of the exogenous genomes, taxonomy trees were constructed per sample, and also for the aggregate samples using Genboree^41^, Fig. 4a. The majority of abundances were assigned to bacteria (89.5%), and 6.4% to Eukaryota, where in terms of majority assignments at the next level, Bacteroidetes/Chlorobi group (28.5%), 27.2% were assigned to Proteobacteria and 17.1% to Firmicutes. Clustering of the overall top taxa by normalized read count was indicative of consistency across samples, with no significant sub-grouping at this level, Fig. 4b.

**Figure 4.**
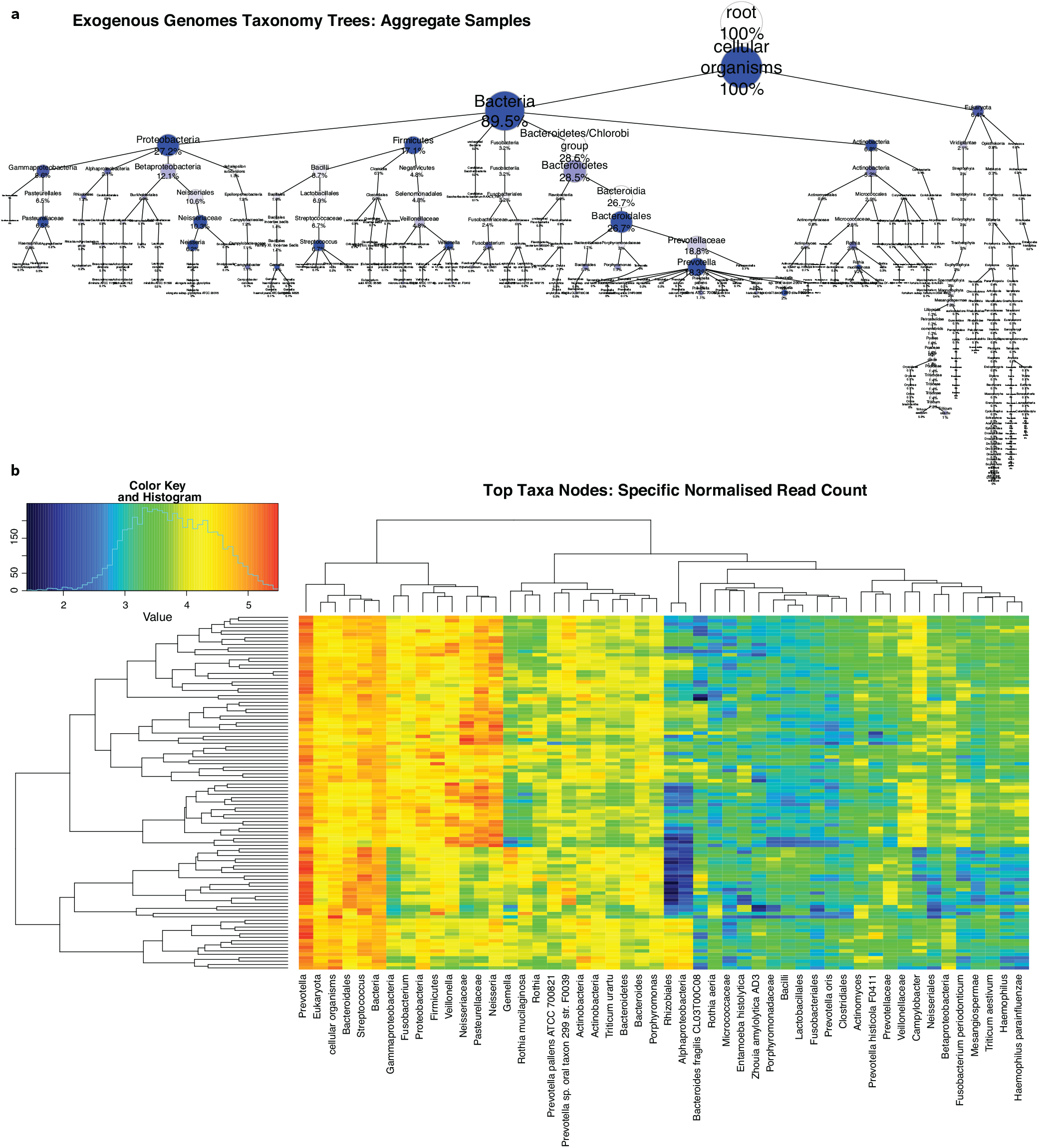
Exogenous taxa from extracellular vesicle analysis. **a** The exogenous genomes taxonomy tree based on mapped EV reads from the aggregate samples is shown. **b** The top taxa nodes indicate consistency across samples (vertical axis).

We again carried out downstream analysis in `MathIOmica`^36^, to create EV time series for mapped data for each of the three time frames (TFH1, TFH2, TFD), and the paired difference (TFΔ) for GENCODE mapped entities, host piRNA, host miRNA and exogenous rRNA and exogenous taxa. Time series were constructed for entities where 0 count values were tagged as Missing, counts < 1 were considered equivalent noise, transcripts with more than 1/4 timepoints missing were removed, and transcripts with constant values across all timepoints were also removed.

### Temporal Trends Identified in Saliva

#### Time Series Classification

The time series for all omics discussed below were classified into temporal trends using `MathIOmica`’s^36^ spectral methods that classify signals based on their autocorrelations, i.e. correlating a time signal with a delayed version of itself, where the delay is characterized as a time *lag* (e.g. lag 1 corresponds to a delay of 1 time interval unit). The method uses a Lomb-Scargle transformation^42–44^ to generate periodograms whose inverse Fourier transform can then produce a set of autocorrelations at different lags for a given time series. Our classification successively identifies time series from the dataset that have statistically significant autocorrelations at particular time lags. In summary (see Methods), `MathIOmica`’s classification generates three sets of classes, strictly based on temporal behavior: 1) Significant autocorrelations at various lags, 2) No autocorrelations, but with positive spikes (abnormally high signals above baselines present at single timepoints), 3) No autocorrelations, but negative spikes present (abnormally low signals below baseline at single timepoints). Within each class a two-tier classification into groups and subgroups is carried out: This approach first separates within-class autocorrelation groups by clustering on autocorrelation lags: signals that may have statistically significant autocorrelation for the class lag, but may still exhibit underlying different structure at other lags. Additionally, the second level clustering into subgroups is based on intensities, and allows us to separate signals that may have different phase (directionality/sense), which cannot be obtained from the periodograms.

The analysis was carried out for each of the omics individually and thousands of individual component trends were identified in the different classes and subgroupings therein. A brief summary is provided in Table 1. The entirety of visualizations and classification memberships are available in the online data files (ODFs), including heatmaps per omic per individual time frame trends, as well as all the code to generate these. We also combined all the classified information to obtain an integrated view of the various omics. Below we showcase parts of the mRNA analysis, as well the results of all the omics combined.

**Table 1.**
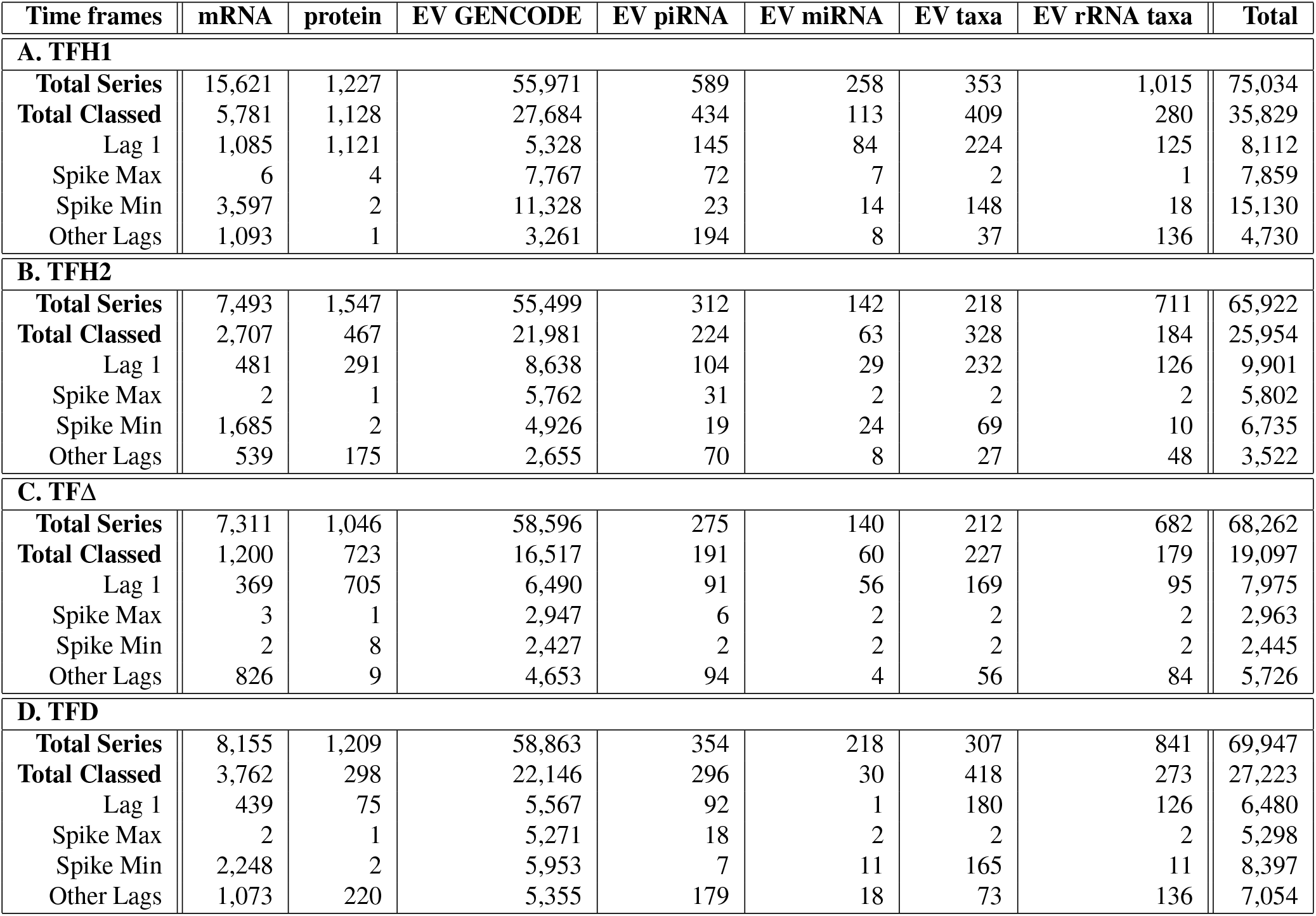
Time series counts and classifications across saliva omics for the various time frames.

#### Saliva mRNA Data Analysis

The trends shown in Fig. 5 correspond to the mRNA time series showing statistically significant time series trends (p-value < 0.01 based on bootstrap simulations, n=100,000) for each of the time frames, for Lag 1 classes.

**Figure 5.**
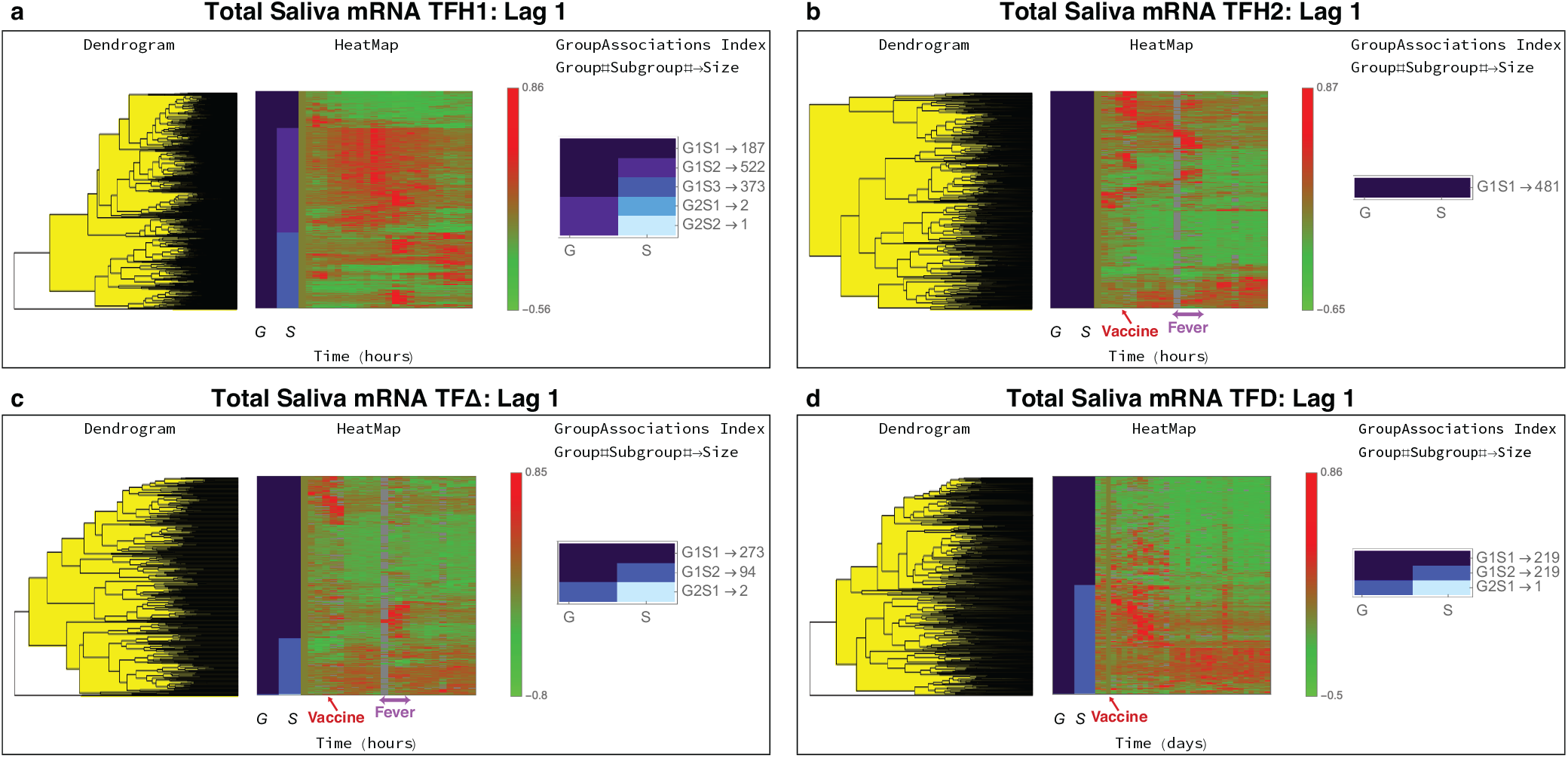
Total saliva mRNA time series trends. The Lag 1 classification results from `MathIOmica` are shown for the different time frames. **a** During TFH1 the subject followed their normal routine. **b** During TFH2, the subject was vaccinated with PPSV23. For both TFH1 and TFH2 the first timepoint corresponds to 7am. Vaccination took place at 10.30 am. The subject reported fever from 5.30pm to 10pm. **c** The TFΔ results correspond to paired differences between TFH2 and TF1 hourly points, to remove intra-day variation so as to focus on the perturbation vaccination responses. The plot is indicative of a response to the vaccine and a response that coincides with the reported fever. **d** For the daily data, TFD, the corresponding vaccine timepoint is Day 3. There is a direct response the days following the vaccination, and a different response in a subset of genes approximately a week following the vaccination, corresponding to immune system activation.

#### Hourly Results (TFH1, TFH2 and TFΔ)

The saliva mRNA showed variation across the day in the untreated TFH1 period. Overall 5,781 time series of mRNA isoforms were found to have statistically significant trends, with 1,085 Lag 1, 6 Spike Max, 3,579 Spike Min and 1,093 other Lag class memberships. The Lag 1 group is shown in Fig. 5a, where the 1,085 time series are further assigned into groups and subgroups based on clustering, according to their different temporal behaviors as described above. In the Lag 1 classification in Fig. 5a there are two groups (G1 and G2). G1 has 3 subgroups (S), S1, S2 and S3 with 187, 522 and 373 time series in each respectively. G2 has 2 subgroups S1, S2 with two and one time series respectively. The groupings show substantial variation in these isoforms’ intensities during the day, with G1S1 showing gradual decreases, G1S2 peaking after morning until the evening, and G1S3 showing peaks later in the evening and night (the first timepoint is at 7am).

In the analysis of the 24 hr period spanning vaccination, TFH2, we should note that the subject reported fever ~7 hrs post vaccination, lasting for about 4.5 hrs. The classification identified 2,707 isoform time series, with 1,685 in Spike Min, 2 in Spike Max, 481 in Lag 1, and 539 in other Lag (≥2) classes. The clustering results for Lag 1 are also shown in Fig. 5b. The changes are indicative of the activation response due to the vaccination.

Given the variation observed in TFH1, we constructed the TFHΔ time series, using paired differences of intensities at each timepoint. The approach aimed to to remove non-vaccination daily variation, and resulted in 1,200 time series with statistically significant trends. These included 369 Lag 1, 3 Spike Max, 2 Spike Min, and 826 other Lags memberships. The Lag 1 results are shown in Fig. 5c. The subgroupings of 273 G1S1 isoform time series, show punctuated trends following vaccination and also coincidental with the reported fever, lasting about 4 hours (timepoints). Furthermore, the G1S2 subgroup of 94 time series is indicative of up-regulation following the vaccination. Additionally a distinct up-regulation of a subset of genes is observed to coincide with the reported fever, Fig. 5c.

We carried out Gene Ontology (GO)^46^ and Reactome Pathway enrichment analysis^45, 47^ and identified multiple involved pathways. Results for TFΔ with False Discovery Rate, FDR < 3 × 10^−3^ are shown in Table 2, and full results available in the ODFs. For the set of TFΔ genes showing immediate response post vaccination and response during the fever period (Class Lag 1, G1S1 Fig. 5c, results include (Table 2A (i)) antigen presentation (class 1 MHC) and processing, interferon alpha/beta and gamma, neutrophil degranulation and ER-Phagosome pathways indicative of the immune activation, including HSF1-dependent transactivation. Furthermore, a set of genes that show continued up-regulation following the vaccination (Class Lag 1, G1S2 Fig. 5c) had enrichment of various immunological pathways including TCR signaling-related pathways, PD-1 signaling, also Interferon gamma signaling, Costimulation by the CD28 family, MHC class II antigen presentation, Cytokine Signaling in Immune system, and also Neutrophil degranulation pathways.

**Table 2.**
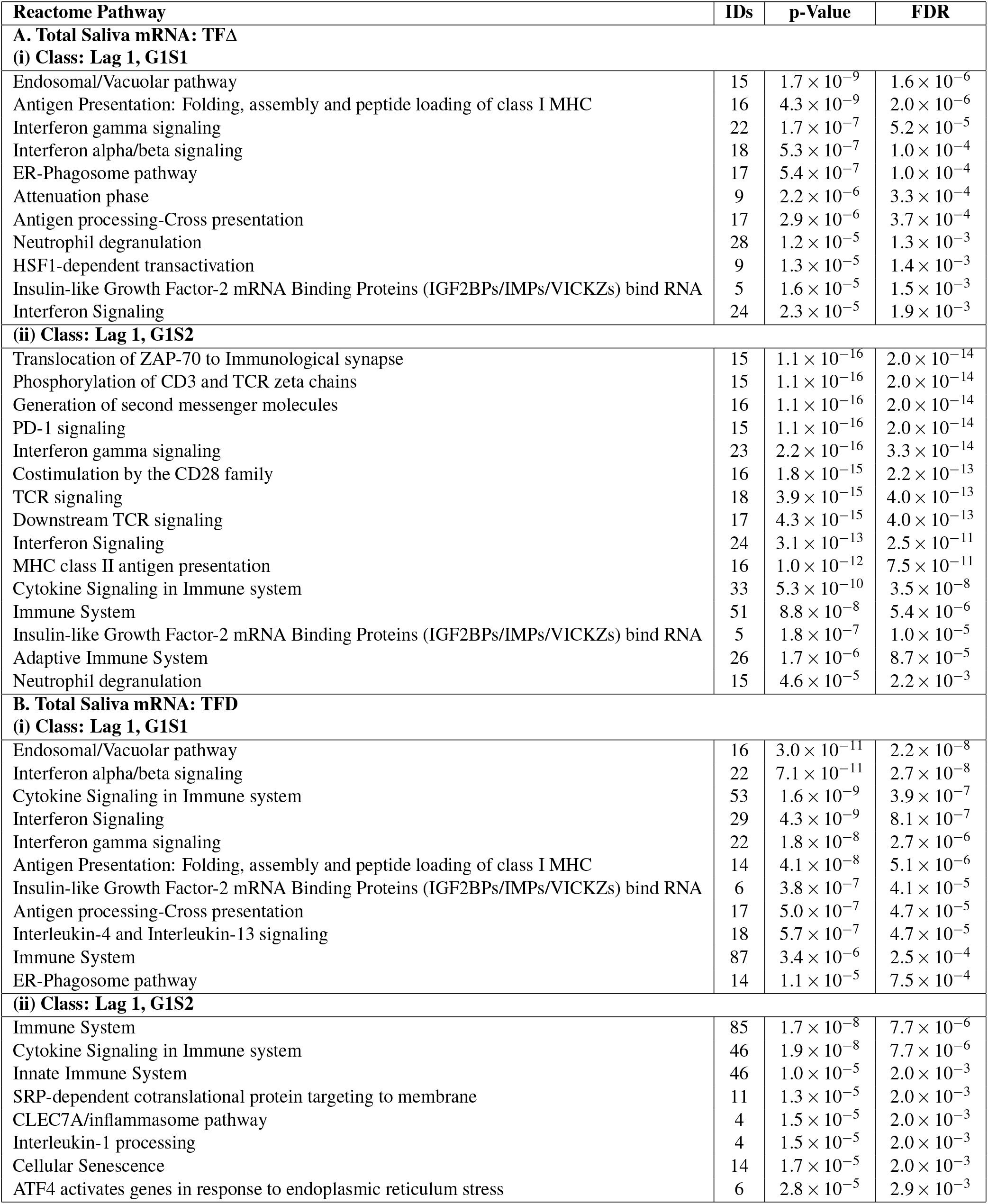
Statistically significant Reactome^45^ pathways identified (FDR< 3 × 10^−3^ results shown).

| Reactome Pathway | IDs | p-Value | FDR |
| --- | --- | --- | --- |
| **A. Total Saliva mRNA: TFA** |  |  |  |
| **(i) Class: Lag 1, G1S1** |  |  |  |
| Endosomal/Vacuolar pathway | 15 | $1.7 \times 10^{-9}$ | $1.6 \times 10^{-6}$ |
| Antigen Presentation: Folding, assembly and peptide loading of class I MHC | 16 | $4.3 \times 10^{-9}$ | $2.0 \times 10^{-6}$ |
| Interferon gamma signaling | 22 | $1.7 \times 10^{-7}$ | $5.2 \times 10^{-5}$ |
| Interferon alpha/beta signaling | 18 | $5.3 \times 10^{-7}$ | $1.0 \times 10^{-4}$ |
| ER-Phagosome pathway | 17 | $5.4 \times 10^{-7}$ | $1.0 \times 10^{-4}$ |
| Attenuation phase | 9 | $2.2 \times 10^{-6}$ | $3.3 \times 10^{-4}$ |
| Antigen processing-Cross presentation | 17 | $2.9 \times 10^{-6}$ | $3.7 \times 10^{-4}$ |
| Neutrophil degranulation | 28 | $1.2 \times 10^{-5}$ | $1.3 \times 10^{-3}$ |
| HSF1-dependent transactivation | 9 | $1.3 \times 10^{-5}$ | $1.4 \times 10^{-3}$ |
| Insulin-like Growth Factor-2 mRNA Binding Proteins (IGF2BPs/IMPs/VICKZs) bind RNA | 5 | $1.6 \times 10^{-5}$ | $1.5 \times 10^{-3}$ |
| Interferon Signaling | 24 | $2.3 \times 10^{-5}$ | $1.9 \times 10^{-3}$ |
| **(ii) Class: Lag 1, G1S2** |  |  |  |
| Translocation of ZAP-70 to Immunological synapse | 15 | $1.1 \times 10^{-16}$ | $2.0 \times 10^{-14}$ |
| Phosphorylation of CD3 and TCR zeta chains | 15 | $1.1 \times 10^{-16}$ | $2.0 \times 10^{-14}$ |
| Generation of second messenger molecules | 16 | $1.1 \times 10^{-16}$ | $2.0 \times 10^{-14}$ |
| PD-1 signaling | 15 | $1.1 \times 10^{-16}$ | $2.0 \times 10^{-14}$ |
| Interferon gamma signaling | 23 | $2.2 \times 10^{-16}$ | $3.3 \times 10^{-14}$ |
| Costimulation by the CD28 family | 16 | $1.8 \times 10^{-15}$ | $2.2 \times 10^{-13}$ |
| TCR signaling | 18 | $3.9 \times 10^{-15}$ | $4.0 \times 10^{-13}$ |
| Downstream TCR signaling | 17 | $4.3 \times 10^{-15}$ | $4.0 \times 10^{-13}$ |
| Interferon Signaling | 24 | $3.1 \times 10^{-13}$ | $2.5 \times 10^{-11}$ |
| MHC class II antigen presentation | 16 | $1.0 \times 10^{-12}$ | $7.5 \times 10^{-11}$ |
| Cytokine Signaling in Immune system | 33 | $5.3 \times 10^{-10}$ | $3.5 \times 10^{-8}$ |
| Immune System | 51 | $8.8 \times 10^{-8}$ | $5.4 \times 10^{-6}$ |
| Insulin-like Growth Factor-2 mRNA Binding Proteins (IGF2BPs/IMPs/VICKZs) bind RNA | 5 | $1.8 \times 10^{-7}$ | $1.0 \times 10^{-5}$ |
| Adaptive Immune System | 26 | $1.7 \times 10^{-6}$ | $8.7 \times 10^{-5}$ |
| Neutrophil degranulation | 15 | $4.6 \times 10^{-5}$ | $2.2 \times 10^{-3}$ |
| **B. Total Saliva mRNA: TFD** |  |  |  |
| **(i) Class: Lag 1, G1S1** |  |  |  |
| Endosomal/Vacuolar pathway | 16 | $3.0 \times 10^{-11}$ | $2.2 \times 10^{-8}$ |
| Interferon alpha/beta signaling | 22 | $7.1 \times 10^{-11}$ | $2.7 \times 10^{-8}$ |
| Cytokine Signaling in Immune system | 53 | $1.6 \times 10^{-9}$ | $3.9 \times 10^{-7}$ |
| Interferon Signaling | 29 | $4.3 \times 10^{-9}$ | $8.1 \times 10^{-7}$ |
| Interferon gamma signaling | 22 | $1.8 \times 10^{-8}$ | $2.7 \times 10^{-6}$ |
| Antigen Presentation: Folding, assembly and peptide loading of class I MHC | 14 | $4.1 \times 10^{-8}$ | $5.1 \times 10^{-6}$ |
| Insulin-like Growth Factor-2 mRNA Binding Proteins (IGF2BPs/IMPs/VICKZs) bind RNA | 6 | $3.8 \times 10^{-7}$ | $4.1 \times 10^{-5}$ |
| Antigen processing-Cross presentation | 17 | $5.0 \times 10^{-7}$ | $4.7 \times 10^{-5}$ |
| Interleukin-4 and Interleukin-13 signaling | 18 | $5.7 \times 10^{-7}$ | $4.7 \times 10^{-5}$ |
| Immune System | 87 | $3.4 \times 10^{-6}$ | $2.5 \times 10^{-4}$ |
| ER-Phagosome pathway | 14 | $1.1 \times 10^{-5}$ | $7.5 \times 10^{-4}$ |
| **(ii) Class: Lag 1, G1S2** |  |  |  |
| Immune System | 85 | $1.7 \times 10^{-8}$ | $7.7 \times 10^{-6}$ |
| Cytokine Signaling in Immune system | 46 | $1.9 \times 10^{-8}$ | $7.7 \times 10^{-6}$ |
| Innate Immune System | 46 | $1.0 \times 10^{-5}$ | $2.0 \times 10^{-3}$ |
| SRP-dependent cotranslational protein targeting to membrane | 11 | $1.3 \times 10^{-5}$ | $2.0 \times 10^{-3}$ |
| CLEC7A/inflammasome pathway | 4 | $1.5 \times 10^{-5}$ | $2.0 \times 10^{-3}$ |
| Interleukin-1 processing | 4 | $1.5 \times 10^{-5}$ | $2.0 \times 10^{-3}$ |
| Cellular Senescence | 14 | $1.7 \times 10^{-5}$ | $2.0 \times 10^{-3}$ |
| ATF4 activates genes in response to endoplasmic reticulum stress | 6 | $2.8 \times 10^{-5}$ | $2.9 \times 10^{-3}$ |

#### Daily Results (TFD)

We also monitored the daily changes post vaccination for 1 month. For the mRNA analysis, 3,762 time series had statistically significant trends. These included 439 Lag 1, 2 Spike Max, 2,248 Spike Min, and 1,073 other Lag memberships. The Lag 1 TFD results are shown in Fig. 5d. As shown in the figure, in the G1S1 subgroup 219 isoform time series show up-regulation, for about 11 days post vaccination, followed by a return to lower expression levels. The 218 time series in the G1S2 subgroup also show a later up-regulation response, after 11 days compared to G1S1, again following the vaccination, and lasting for the remainder of the daily observation period.

Reactome pathway and GO enrichment analysis also identified multiple pathways corresponding to each trend. Reactome results for TFD Lag 1 are shown in Table 2B(i)-(ii) for Lag 1 G1S1 and G1S2 subgroups (full results, including GO terms available in the ODFs). In the TFD Lag 1 G1S1 group, over-representation included Interferon Signaling (29 genes), Cytokine Signaling in Immune system (53 genes), Antigen Presentation (MHC I related) and inteleukin 4 and 13 signaling pathways (18 genes). These are indicative of an early response within the first days after vaccination. For TFD Lag 1 G1S2 the results included general Immune System activation (85 genes), and also Cytokine Signaling in Immune System (46 genes).

### Other Omics

In addition to the mRNA, each other set of omics was individualized analyzed to identify temporal trends, using the same classification method in `MathIOmica` as described above. This identified statistically significant trends for time series for different omics in the different time frames are shown in Table 1.

The different classes for the omics datasets were joined within each respective time frame, and data within each combined class were clustered together. The breakdown of identified trends included overall 35,829 time series for TFH1, 25,954 for TFH2, 19,097 for TFΔ, and 27,223 for TFD. In reference to the corresponding omics, the EV GENCODE identifiers accounted for more that 77% of the time series across all time frames. The results from Lag 1 are again shown in Fig. 6. In terms of temporal behavior, we notice again similar responses in the various time frames. In TFH1, we again see large temporal variation, with sets of time series being up-regulated during awake time and a subset during night time as shown in Fig. 6a. In TFH2, Fig. 6b, the effects of the vaccination become apparent with up-regulated responses following the vaccination.

**Figure 6.**
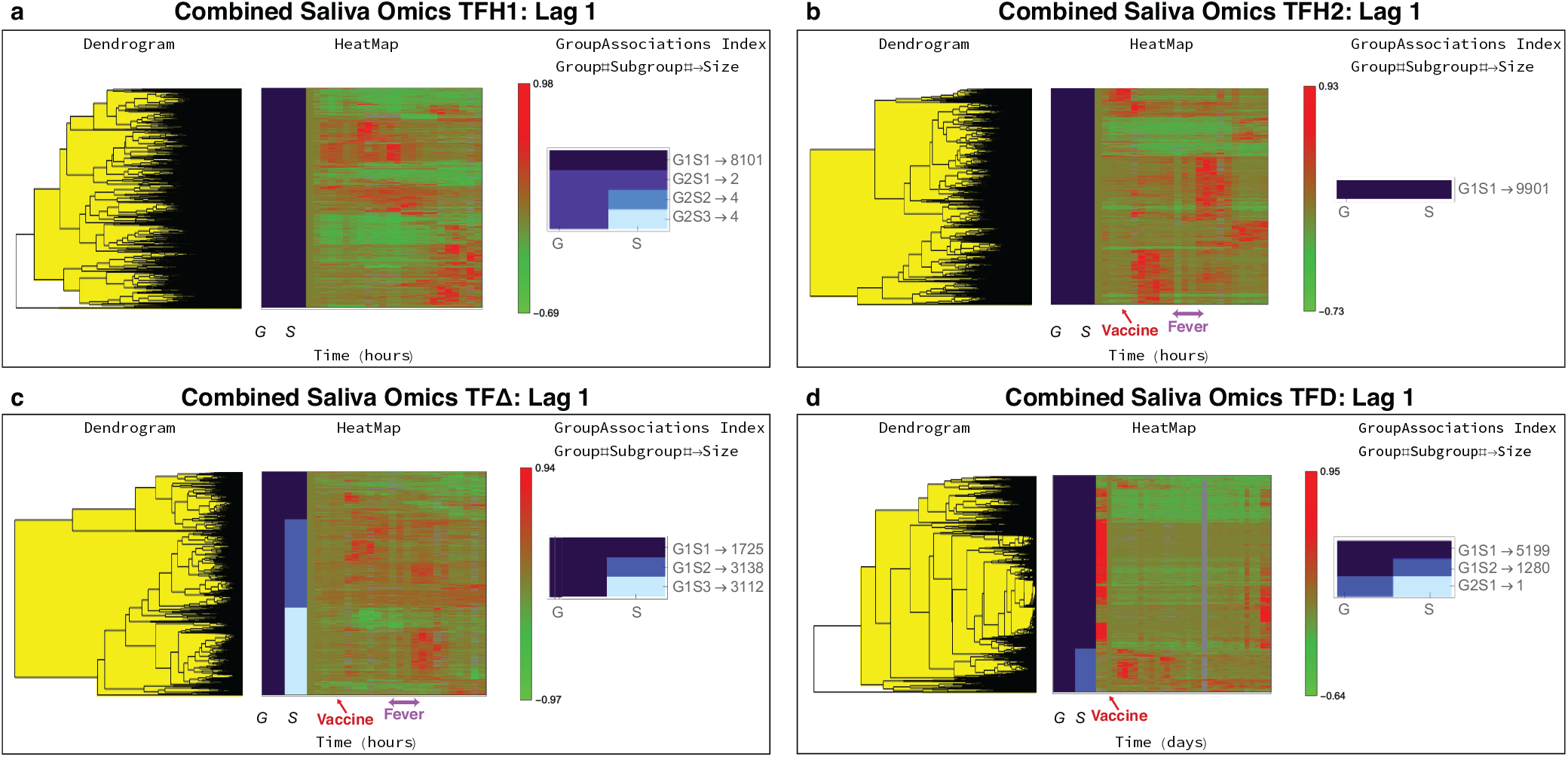
Combined saliva omics time series trends. The Lag 1 classification results from `MathIOmica` are shown for the different time frames. **a** During TFH1 the subject followed their normal routine. **b** During TFH2, the subject was vaccinated with PPSV23. For both TFH1 and TFH2 the first timepoint corresponds to 7 am. Vaccination took place at 10.30 am. The subject reported fever from 5.30 pm to 10 pm. **c** The TFΔ results correspond to paired differences between TFH2 and TFH1 hourly points, to remove intra-day variation so as to focus on the perturbation vaccination responses. The plot is indicative of a phased response to the vaccine compared to the mRNA responses and a response that is again shifted compared to the reported fever (cf. Fig. 5c). **d** For the daily data, TFD, the corresponding vaccine timepoint is Day 3. There is a direct response the days following the vaccination. The EV omics dominate the information in this plot, compared to the mRNA in total saliva response (cf. Fig. 5d).

In the paired comparison TFΔ for Lag 1 in Fig. 6c, there are 3 subgroups, G1S1 with 1,725 time series, showing increase in intensity about 2-3 hours post vaccination and subsequent decrease; G1S2 with 3,138 time series showing increase 2-3 hours post vaccination for a few hours, and additionally again increase in intensity about 14 hours post vaccination (after the fever time span); and G1S3 with 3,112 components with increases in intensity about 14 hours post vaccination (again after the fever time span). We note here that the response/time pattern appear lagging compared to the corresponding mRNA results in Fig. 5c, by approximately 3 hours, both following the immediate vaccine response, and also for the reported fever time span. The mRNA over-representation analysis was discussed above. Proteomics displayed similar trends to the mRNA. For Lag 1 the Reactome analysis for the aggregate group/subgroup proteins resulted in multiple statistically significant pathways (FDR < 0.01), and included Nonsense Mediated Decay Pathways, Neutrophil degranulation, various translation related pathways, Immune System (with 224 entities identified), and more (the top pathways are shown in Table 3A, FDR < 1.2×10^−11^).

**Table 3.**
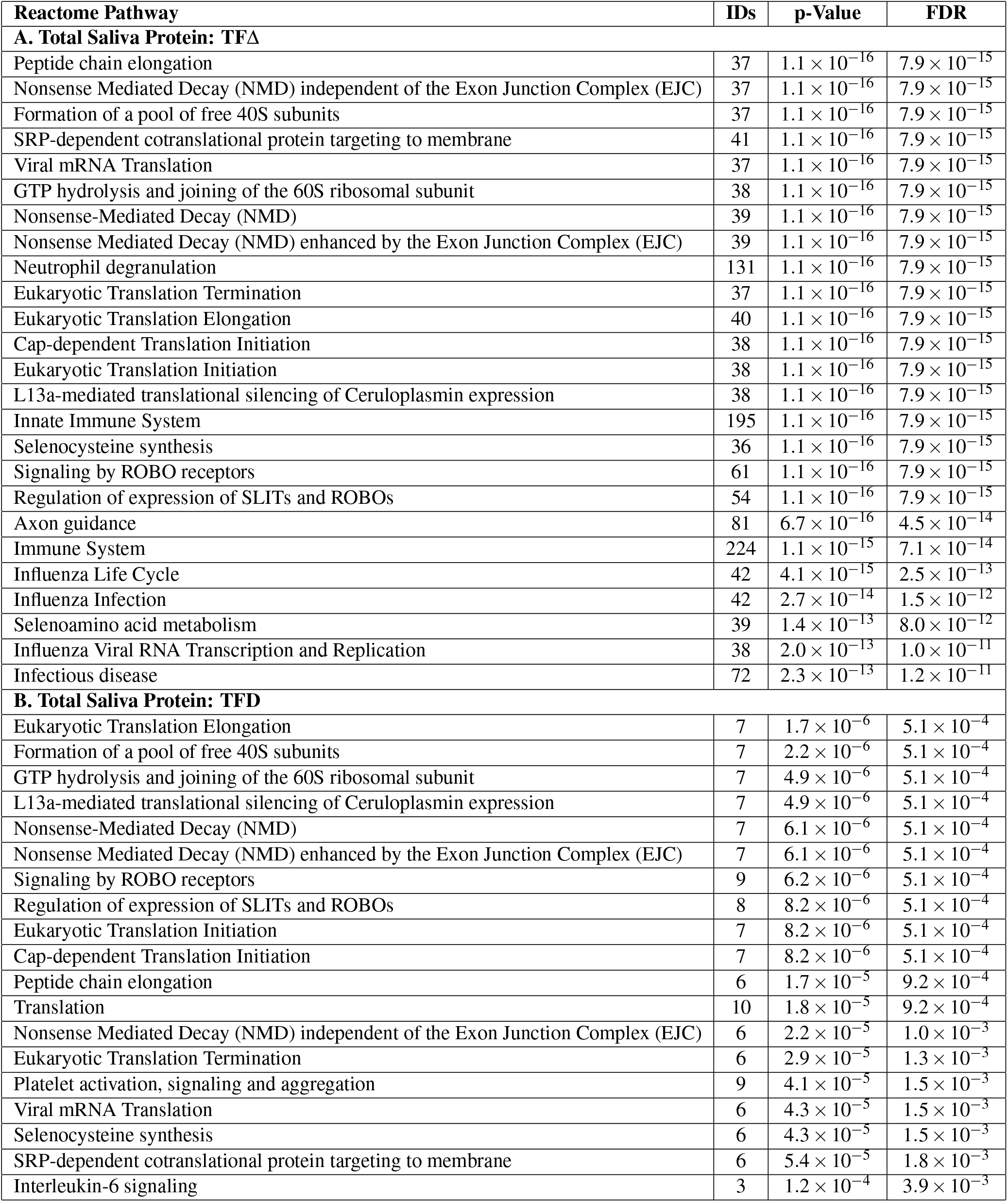
Reactome pathway enrichment analysis results for protein Lag 1 aggregate results for TFΔ and TFD

| Reactome Pathway | IDs | p-Value | FDR |
| --- | --- | --- | --- |
| **A. Total Saliva Protein: TFA** |  |  |  |
| Peptide chain elongation | 37 | $1.1 \times 10^{-16}$ | $7.9 \times 10^{-15}$ |
| Nonsense Mediated Decay (NMD) independent of the Exon Junction Complex (EJC) | 37 | $1.1 \times 10^{-16}$ | $7.9 \times 10^{-15}$ |
| Formation of a pool of free 40S subunits | 37 | $1.1 \times 10^{-16}$ | $7.9 \times 10^{-15}$ |
| SRP-dependent cotranslational protein targeting to membrane | 41 | $1.1 \times 10^{-16}$ | $7.9 \times 10^{-15}$ |
| Viral mRNA Translation | 37 | $1.1 \times 10^{-16}$ | $7.9 \times 10^{-15}$ |
| GTP hydrolysis and joining of the 60S ribosomal subunit | 38 | $1.1 \times 10^{-16}$ | $7.9 \times 10^{-15}$ |
| Nonsense-Mediated Decay (NMD) | 39 | $1.1 \times 10^{-16}$ | $7.9 \times 10^{-15}$ |
| Nonsense Mediated Decay (NMD) enhanced by the Exon Junction Complex (EJC) | 39 | $1.1 \times 10^{-16}$ | $7.9 \times 10^{-15}$ |
| Neutrophil degranulation | 131 | $1.1 \times 10^{-16}$ | $7.9 \times 10^{-15}$ |
| Eukaryotic Translation Termination | 37 | $1.1 \times 10^{-16}$ | $7.9 \times 10^{-15}$ |
| Eukaryotic Translation Elongation | 40 | $1.1 \times 10^{-16}$ | $7.9 \times 10^{-15}$ |
| Cap-dependent Translation Initiation | 38 | $1.1 \times 10^{-16}$ | $7.9 \times 10^{-15}$ |
| Eukaryotic Translation Initiation | 38 | $1.1 \times 10^{-16}$ | $7.9 \times 10^{-15}$ |
| L13a-mediated translational silencing of Ceruloplasmin expression | 38 | $1.1 \times 10^{-16}$ | $7.9 \times 10^{-15}$ |
| Innate Immune System | 195 | $1.1 \times 10^{-16}$ | $7.9 \times 10^{-15}$ |
| Selenocysteine synthesis | 36 | $1.1 \times 10^{-16}$ | $7.9 \times 10^{-15}$ |
| Signaling by ROBO receptors | 61 | $1.1 \times 10^{-16}$ | $7.9 \times 10^{-15}$ |
| Regulation of expression of SLITs and ROBOs | 54 | $1.1 \times 10^{-16}$ | $7.9 \times 10^{-15}$ |
| Axon guidance | 81 | $6.7 \times 10^{-16}$ | $4.5 \times 10^{-14}$ |
| Immune System | 224 | $1.1 \times 10^{-15}$ | $7.1 \times 10^{-14}$ |
| Influenza Life Cycle | 42 | $4.1 \times 10^{-15}$ | $2.5 \times 10^{-13}$ |
| Influenza Infection | 42 | $2.7 \times 10^{-14}$ | $1.5 \times 10^{-12}$ |
| Selenoamino acid metabolism | 39 | $1.4 \times 10^{-13}$ | $8.0 \times 10^{-12}$ |
| Influenza Viral RNA Transcription and Replication | 38 | $2.0 \times 10^{-13}$ | $1.0 \times 10^{-11}$ |
| Infectious disease | 72 | $2.3 \times 10^{-13}$ | $1.2 \times 10^{-11}$ |
| **B. Total Saliva Protein: TFD** |  |  |  |
| Eukaryotic Translation Elongation | 7 | $1.7 \times 10^{-6}$ | $5.1 \times 10^{-4}$ |
| Formation of a pool of free 40S subunits | 7 | $2.2 \times 10^{-6}$ | $5.1 \times 10^{-4}$ |
| GTP hydrolysis and joining of the 60S ribosomal subunit | 7 | $4.9 \times 10^{-6}$ | $5.1 \times 10^{-4}$ |
| L13a-mediated translational silencing of Ceruloplasmin expression | 7 | $4.9 \times 10^{-6}$ | $5.1 \times 10^{-4}$ |
| Nonsense-Mediated Decay (NMD) | 7 | $6.1 \times 10^{-6}$ | $5.1 \times 10^{-4}$ |
| Nonsense Mediated Decay (NMD) enhanced by the Exon Junction Complex (EJC) | 7 | $6.1 \times 10^{-6}$ | $5.1 \times 10^{-4}$ |
| Signaling by ROBO receptors | 9 | $6.2 \times 10^{-6}$ | $5.1 \times 10^{-4}$ |
| Regulation of expression of SLITs and ROBOs | 8 | $8.2 \times 10^{-6}$ | $5.1 \times 10^{-4}$ |
| Eukaryotic Translation Initiation | 7 | $8.2 \times 10^{-6}$ | $5.1 \times 10^{-4}$ |
| Cap-dependent Translation Initiation | 7 | $8.2 \times 10^{-6}$ | $5.1 \times 10^{-4}$ |
| Peptide chain elongation | 6 | $1.7 \times 10^{-5}$ | $9.2 \times 10^{-4}$ |
| Translation | 10 | $1.8 \times 10^{-5}$ | $9.2 \times 10^{-4}$ |
| Nonsense Mediated Decay (NMD) independent of the Exon Junction Complex (EJC) | 6 | $2.2 \times 10^{-5}$ | $1.0 \times 10^{-3}$ |
| Eukaryotic Translation Termination | 6 | $2.9 \times 10^{-5}$ | $1.3 \times 10^{-3}$ |
| Platelet activation, signaling and aggregation | 9 | $4.1 \times 10^{-5}$ | $1.5 \times 10^{-3}$ |
| Viral mRNA Translation | 6 | $4.3 \times 10^{-5}$ | $1.5 \times 10^{-3}$ |
| Selenocysteine synthesis | 6 | $4.3 \times 10^{-5}$ | $1.5 \times 10^{-3}$ |
| SRP-dependent cotranslational protein targeting to membrane | 6 | $5.4 \times 10^{-5}$ | $1.8 \times 10^{-3}$ |
| Interleukin-6 signaling | 3 | $1.2 \times 10^{-4}$ | $3.9 \times 10^{-3}$ |

GO analysis for the EV GENCODE results for TFΔ Lag 1 were generic and included multiple cellular processes, and even though innate immunity (56 IDs, adjusted p-value < 6.0×10^−4^) and adaptive immunity (21 IDs, adjusted p-value < 0.01) were also detected as enriched in the aggregate (all subgroups) analysis. The results appear as non-specific to immune response, particularly as no over-representation in Reactome pathways was found to be statistically significant based on FDR. For EV miRNA the TFΔ Lag 1 showed statistically significant over-representation in GTP binding (p-value < 4.6×10^−14^, perinuclear region of cytoplasm (p-value < 9.3×10^−14^, nerve growth factor receptor signaling pathway 1.5×^10−13^, MAPK cascade 1.7×10^−13^, MYD88 dependent toll-like receptor signaling pathway 1.7×10^−13^, toll-like receptor signaling pathway 3.4×10^−13^ and others.

Furthermore, in the TFD combined omics results, there are two main trends, Fig. 5d, with G1S1 showing 5,199 series showing a relative decrease in intensity, for which some return to pre-immunization levels after about one month, and G1S2 with 1,280 components showing a set of time series that increase in intensity immediately post vaccination for approximately 12 days, and a set that remains high in intensity for the duration of the month’s measurements. The mRNA TFD pathway analysis results were discussed above, Table2B. Proteomics time series on the other hand displayed fewer pathway enrichment results as shown in Table 3B, including various Nonsense-Mediated Decay pathways, Translation related pathways, L13a-mediated translational silencing of Ceruloplasmin expression, Platelet activation and others. GO analysis for the EV GENCODE results for TFD Lag 1 included biological processes relating to transcription and its regulation, and also included innate immunity (65 IDs, adjusted p-value < 2.8×10^−5^). Full enrichment analysis results for all classifications and omics is available in the ODFs on Zenodo.

To identify a set of time series omics that showed statistically significant temporal trends (adjusted p-value < 0.01) in multiple time frames, we intersected the results for Lag 1 for the various omics for Lag 1 TFΔ and TFD time frames. The overlaps for the omics considered included 43 mRNA, 44 protein, 658 EV GENCODE, 30 EV piRNA, 41 EV exogenous taxa, and 45 EV rRNA taxa time series (mRNA, protein, piRNA and exogenous taxa memberships are shown in Table 4). We carried out Reactome pathway analysis on the overlaps directly. For mRNA we observed over-representation in Insulin-like Growth Factor-2 mRNA Binding Proteins (IGF2BPs/IMPs/VICKZs) bind RNA (FDR < 5.9×10^−6^, CLEC7A/inflammasome pathway (FDR < 3.0×10^−3^ Interleukin-4 and Interleukin-13 signaling (FDR < 0.026) - the genes involved include CD44 and IL1B, that are associated with monocyte aggregation, and FCGR2B and SAMSN1 that are involved in negative regulation of B-cell proliferation. For proteins, pathway associations include Signaling by ROBO receptors (FDR < 2.8×10^−3^), Eukaryotic Translation Elongation (FDR < 2.8×10^−3^), Formation of a pool of free 40S subunits (FDR < 2.8×10^−3^), Interleukin-6 signaling FDR (FDR < 2.8×10^−3^) and Neutrophil degranulation (FDR < 4.4×10^−3^). Other omics overlaps did not show statistically significant over-representations.

**Table 4.**
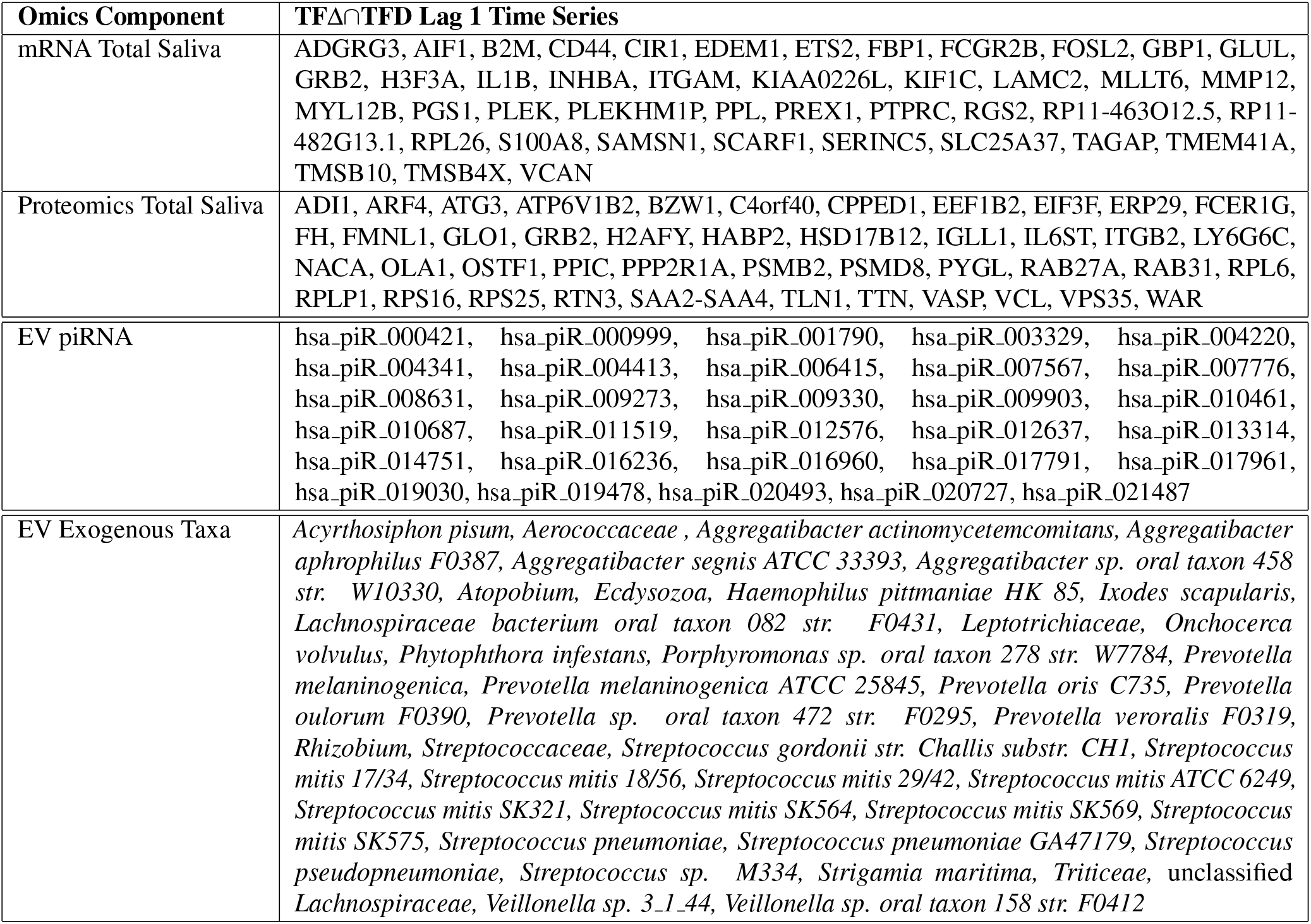
Overlaps of responding components between time frames TFΔ and TFD.

Finally, for each class of the combined classified data, we constructed putative interaction networks based on the Euclidean distance between the class members. Here, we assumed that the omics showing similar trends over time within each class are likely to be associated to each other (even though interactions may be indirect). The group/subgroup annotations within each class were also included. The constructed networks are available as a resource in the ODFs, including *.json* files, and may be used to explore and validate possible interactions in saliva data (see online Methods).

## Discussion

We have presented here our findings from a case study of the utility of saliva towards personalized health monitoring. Following vaccination of a subject with pneumococcal vaccine (PPSV23) we were able to detect distinct signatures in various saliva omics. We were able to profile more than 65,000 components in various time frames over time, and identify 19,000+ time series that had statistically significant temporal trends. The time series trends observed were indicative of immune response, which coincided in timing with the vaccination, and fever reported by our subject. The time frames of immune responses observed are concordant with our expectations of innate and adaptive immunity development, as seen both in the immediate hourly as well as the short- and long-term daily responses observed. Various pathways were activated, involved in immune response and regulation, including interferon signaling and MHC antigen presentation. The immune activation spanned an initial response within hours, as well as long term response extending for over a month.

Our results suggest that saliva omics can be consistently assessed for personalized monitoring. While multiple omics provide responses post-vaccination as discussed for the TFΔ and TFD time frames, mRNA results appear more specific and sensitive (time-wise) to the vaccination. EV response, particularly transcript-level (GENCODE), though very sensitive and responsive does not yield as specific results as the mRNA from total saliva. In fact too many EV mRNAs show statistically significant temporal responses, which is likely due to increased EV release from various cells, but not specific in terms of reflecting the functional responses in the cells of origin. EV results also suggest a lagged response by a few hours compared to the mRNA observations, which also suggests that the mRNA measurements can potentially provide more timely data for practical health monitoring. Additional omics showed responses concordant with the mRNA responses, including miRNA, piRNA and exogenous taxa quantitations, however these need further validation, particularly as our knowledgebase of pathway implications and functional important of such omics is still under development. Some of omics time series are found in responses across hourly and daily samplings, Table 4, and such sets of omics can be targeted for non-invasive health monitoring. The processing of samples for mRNA involved the use of standardized kits that can be used by subjects remotely, and can facilitate storage of sample for about a month without refrigeration. Additionally, the mRNA sequencing preparation and result analysis is considerably faster than the other omics processing, so our recommendation based on our findings is to utilize similar approaches for mRNA broad profiling, while using targeted protein/EV-content assays. These omics must be coupled with standard physiological and molecular measures already utilized in the clinic for a complete assessment of health status. While our results are based on PPSV23 activation, we anticipate that they can be extended to additional vaccine and immune disease profiling, particularly with the goal of discovering immune-specific signatures for each affliction and/or intervention.

Our study has limitations. Even though we attempted to pair time responses for the hourly data, this is still a single subject case-study (n=1), and our results will need to be validated. Furthermore, due to limited samples and resources, we did not carry out immune profiling, such as cytokine assays or functional assays to assess the immune acquisition to the different components in the PPSV23 vaccine. We were also unable to obtain blood samples across all the time points, as our focus was a first evaluation of saliva omics. Additionally, our study did not specifically target salivary microbiome, and the meta-analysis of EV RNA content indicated substantial variability in overall taxa, the composition of which is expected to vary across individuals. In the next stage of our long-term project we have collected samples from multiple subjects being vaccinated at the same time points with PPSV23 and monitored over multiple timepoints. Given further resources, we plan to utilize these samples to both validate and extend our findings to also include monitoring of blood components. By comparing the responses in blood and saliva we will be able to assess to what extend saliva may be used as a proxy for blood monitoring, identifying common and different responses in different tissues. Finally, in monitoring total saliva omics in bulk, we are ignoring the multi-cellular composition of saliva. With the availability of single-cell RNA-seq methodology, we anticipate that we will be able to also assess the cell-type-specific response in saliva.

In summary, saliva provides a promising venue for non-invasive diagnostics of immune response. This is particularly important for enhancing our diagnostic capabilities for multiple viral or bacterial responses, particularly in cases where blood may not be easily available, due to technical issues, remote locations (e.g. monitoring active personnel), lack of specialized equipment and healthcare availability (e.g. due to socio-economic factors), patient vulnerability (immuno-compromised, children, and elderly populations). Given the current pandemic (COVID-19), enhancing our diagnostic capabilities has become a high priority. While the utility of saliva for differentiating between different afflictions still needs to be evaluated, our study provides the first steps towards a no-pain no-blood diagnostic process that can greatly enhance our capabilities for universal individualized health care and diagnostics.

## Methods

### Ethical Approval

All experimental protocols were approved by the Institutional Review Board under protocol number LEGACY15-071 (15-071) at Michigan State University. All methods were carried out in accordance with the relevant guidelines and regulations.

### Data and Protocol Availability

Sequencing data reported here are available on Gene Expression Omnibus under accessions GSE108664 (Saliva mRNA-sequencing) and GSE108666 (EV small RNA). Proteomics data are available on MassIVE as part of accession MSV000081869. All scripts and data analysis code utilized in the integrative analysis are available on Zenodo (DOI:10.5281/zenodo.3358011) as online data files (ODFs), in addition to results and methods as referred to in the manuscript.

### Sample Collection

Samples were taken in the three time frames hourly for TFH1 and TFH2, and daily for TFD. In TFH2, the PPSV23 pneumococcal vaccine was administered at approximately 10.30 am (between sample collections at 10 and 11 am). Following the vaccination, and after the 24 hour monitoring, daily samples were taken for about a month. At each timepoint 5 ml saliva were collected: 2 ml in an Oragene (DNAGenotek) tube for RNA sequencing, and 3 ml in a conical tube for EV characterization and mass spectrometry proteomics, as described below. Daily samples were all taken at 8 am, to limit variability. Additionally the subject followed the exact same diet and meal timings during TFH1 and TFH2, and had neither meals/drinks nor teeth brushing for at least 1 hour prior to TFD sample donations.

### Saliva Sample Processing for RNA-Sequencing

The saliva samples (2 ml) for RNA processing were collected in Oragene (DNAGenotek) tubes. The samples were incubated at 50°C for 1 hour, and stored at −80 °C. RNA Processing: 500 *μ*l aliquots were incubated at 90 °C for 15 minutes in a heating block and then then cooled to room temperature. 20 *μ*l neutralizer solution were mixed with each aliquot, vortexed and incubated on ice for 10 minutes, precipitating impurities and inhibitors. The sample was then centrifuged at 13,000 g for 3 minutes. The supernatant was transferred into a new microcentrifuge tube, 2 volumes of cold 95% EtOH were added and mixed thoroughly, followed by incubation at −20 °C for 30 minutes. Following centrifugation at 13,000 g for 3 minutes, the precipitate was collected. This pellet was dissolved in 350 *μ*l of buffer RLT (RNeasy Micro kit). 350 *μ*l of 70% ethanol were added and mixed. Additional steps followed the RNeasy Cleanup (Qiagen) per manufacturer’s instructions to obtain concentrated RNA.

Libraries were constructed and sequenced by Novogene, using a Eukaryotic directional mRNA library (NEB). cDNA preliminary concentration was quantitated on a Qubit (Life Technologies), an Agilent Bioanalyzer 2100 was used to test the insert size, and Q-PCR was used to quantify the library effective concentration precisely. The cDNA libraries were sequenced on an Illumina HiSeq 4000.

### Saliva EV Processing

Saliva samples for EV processing were collected in a conical tube and stored in −80°C on sample receipt. EVs were processed from 500 *μ*l saliva, following centrifugation at 3000 g for 20 minutes at 4 °C. 500 *μ*l of saliva were centrifuged at 3000 g for 20 minutes at 4 °C to remove cells and cell debris. ExoQuick-TC Exosome Precipitation Solution (SBI) was added to the supernatant in a 2:1 ratio, and the mixture was refrigerated overnight at 4 °C. Following incubation samples were centrifuged 1500g for 30 minutes at 4 °C. The supernatant was aspirated and residual ExoQuick-TC solution was spun down at 1500 g for 5 minutes. EV pellets were stored at −80 °C. RNA was extracted using the SeraMir RNA Amplification Kit (SBI) per manufacturer’s instructions. The quality of EV RNA was checked using a 2100 Bioanalyzer (Agilent).

Small RNA library preparation was carried out using NEBNext Multiplex Small RNA library prep kit (New England Biolabs) following manufacturer’s instructions. After PCR amplification, quality of libraries was assessed using a high sensitivity DNA kit on a Bioanalyzer (Agilent) according to manufacturer’s instructions. Size selection was performed using 3% agarose dye-free marker H cassettes on a Pippin Prep (Sage Science) following manufacturer’s instructions with a specified collection size range of 125–153 bp. Libraries were further purified and concentrated by ethanol precipitation, resuspended in 10 *μ*l of 10 mM tris-HCl (pH=8.5) and quantified using Qubit and Bioanalyzer. Based on the quantification, equimolar library pools were prepared, quality was assessed as described above and the library was further diluted to 4 nM using 10 mM tris-HCl (pH=8.5). Pooled libraries were sequenced at a final concentration of 1.2 pM on an Illumina HiSeq 2500 (15-plex, 1×50 bp format).

### Exosome Quantitation by ELISA

EV concentrations were quantitated using the EXOCET Exosome Quantitation Assay kit (SBI). EVs from 1ml of saliva were precipitated using the Exoquick TC protocol (see above). Each exosome pellet was dissolved in 80 *μ*l of lysis buffer and diluted with 80 *μ*l of PBS to be used for duplicate reactions. Samples were then incubated at 37 °C for 5 minutes to liberate EV proteins, vortexed for 15 seconds, and centrifuged at 1500 g for 5 minutes to remove debris. The supernatant EV protein samples were then assayed on a microtiter plate following the EXOCET kit manufacturer protocol (SBI), including 7 standards and blanks in duplicates. The plate was read using a spectrophotometric plate reader (Bio-RAD) at 405 nm. Spectrophotometry results for standards were used to obtain a linear fit, and sample results were indicative of ~10^9^ EVs/ml (supplementary data on Zenodo).

### EV Transmission Electron Microscopy

Isolated EVs were fixed in 2% paraformaldehyde (PFA) for 5 min. For negative-staining of EVs, 5*μ*l of the sample solution was placed on a carbon-coated EM grid and EVs were immobilized for 1 min. Next, the grid was transferred to five 100 *μ*l drops of distilled water and letting it for 2 min on each drop. The sample was negative-stained with 1% uranyl acetate. The excess uranyl acetate was removed by contacting the grid edge with filter paper and the grid was air-dried. The grids were imaged with a JEOL 100CXII Transmission Electron Microscope operating at 100 kV. Images were captured on a Gatan Orius Digital Camera.

### Nanoparticle Tracking Analysis (NTA)

NTA was carried out using the ZetaView (Particle Metrix) following the manufacturer’s instructions. EVs derived from saliva were further diluted 1000- to 5000-fold with PBS for the measurement of particle size and concentration.

### Saliva Proteomics (Mass Spectrometry)

Saliva samples for proteomics processing were collected in a conical tube (same as EVs - see above) and stored in −80 °C on sample receipt. For proteomics processing, the Tandem Mass Tag (TMT) 6-plex kits were used (Thermo). Per sample, 300 *μ*l of saliva were and dissolved in 300 *μ*l lysis buffer (1:1 ratio saliva to lysis buffer to achieve > 2mg/ml protein concentration). Protein concentration were evaluated using a Qbit (Life sciences). Per timepoint, 100 *μ*g were used, adjuste to a final volume of 100 *μ*L with 100 mM TEAB. The manufacturer’s TMT labeling protocol was then followed to prepare the protein extract (part A, steps 7 onwards, and parts B for protein digestion and C for peptide labeling). Samples were ran through OffGel Fractionation (using an Agilent Offgel 3100 fractionator), and mass sectrometry was carried out with a ThermoFisher Q-Exactive mass spectrometer (www.thermo.com) using a FlexSpray nano-spray ion source. Survey scans were taken in the Orbi trap (70,000 resolution, determined at m/z 200) and the top twelve ions in each survey scan are then subjected to automatic higher energy collision induced dissociation (HCD) with fragment spectra acquired at 35,000 resolution. Additional details are provided in the online experimental protocols on Zenodo (see above).

### Data Mapping

#### Total Saliva RNA-seq

Fastq files from paired-end sequencing (150 bp paired-end reads) were mapped using `Kallisto`^31, 32^ (with bootstrap sample parameter, −b, set to 100. For annotation, GENCODE^35^ v28 transcripts and genome built GRCh38.p12 were used. The mapping results across timepoints were compiled using `sleuth`^33^(with `DESeq`^34^ adjustment of Transcripts per Million). We note that the annotation used gene name concatenated with ‘kind’ information (`‘ext gene’:‘kind’`).

The transcriptomics results were imported as `OmicsObject` constructs in `MathIOmica`^36^. Zero intensities were tagged as missing values, and intensities with aTPM < 1 were set to unity. Time series were constructed only for transcripts for which a signal was detected for at least 3/4 of the time points, and also constant time series were removed.

#### Total Saliva Proteomics

Proteomics *.raw* mass spectrometry files were analyzed using Proteome Discoverer (Thermo), using UniProt^37^ human proteome database for reference. Mass tolerance was set to 10 ppm for precursor ions, and to 0.02 Dalton for fragment ions. Modifications included cysteine carbamidomethylation (fixed) and N-terminal and lysine TMT 6-plex and methionine oxidation (variable). Furthermore, we allowed for < 2 trypsin digestion missed cleavages. Proteins were identified using unique peptides of length ≥ 6 amino acids. We set FDR < 1% (strict) and < 5% (non-strict). For identification, we calculated results for both cases with 1 or 2 unique peptides per protein. We carried out peptide quantitation using unique peptides (reporter ion mass tolerance *<* 10 ppm). For protein quantitation, we used medians of peptide ratios.

Multi-consensus reports from each set of technical replicates were constructed and used downstream in `MathIOmica` to construct an annotated `OmicsObject`. For each timepoint (sample) a Box-Cox power transformation was first used to transform the data to normal distributions^38^. Time series were constructed only for proteins with at least 1 unique peptide, and for which a signal was detected for at least 3/4 of the time points. Constant time series were also removed.

#### EVs Small RNA-seq

Small RNA-seq data from EVs were processed using the Genboree Workbench^40, 41, 48^ `exceRpt` pipeline^39^ to assess content by: (1) First removing reads that map to UniVec contaminants, 45S, 5S and mitochondiral rRNAs; (2) mapping reads sequentially to human miRNAs (mirBase), tRNAs (gtRNAdb), piRNAs(piRNABank), GENCODE and circRNAs (cirBase); (3) mapping unmapped reads from (2) to exogenous miRNAs and rRNAs; (4) finally mapping unmapped reads from (3) to all genomes in ensembl and NCBI. Parameter settings and Genboree output files are available on Zenodo (see data availability).

For each biotype `MathIOmica OmicsObject` constructs were created. Zero intensities were tagged as missing values, and intensities with aTPM *<* 1 were set to unity. Time series were constructed only for transcripts for which a signal was detected for at least 3/4 of the time points, and also constant time series were removed.

#### Temporal Analysis and Integration

For all mapped data time series were constructed with reference to the first timepoint for TFH1, TFH2 and TFΔ, and with reference to the vaccination day for TFDaily. TFΔ time series were constructed as paired hourly timepoint intensity differences between TFH2 and TFH1. All series were normalized as vectors to unit length. Time series classification analyses were carried out using `MathIOmica` as detailed below and in the online Mathematica notebooks^36, 49^.

#### Temporal Classification Details

The time series classification used `MathIOmica’s TimeSeriesClassification` function with the `Method −> “Autocorrelation”` setting^36^. Briefly, for a given omic signal *j* with *X_j_* intensities over *N* times we construct a time series *X_j_* = {*X_j_*(*t*_1_)*, X_j_*(*t*_2_) *…, X_j_*(*t_N_*)}. The signal’s periodogram is obtained using a Lomb-Scargle transformation^42–44^ to account for uneven sampling, *P_LS_*. An inverse Fourier transform on *P_LS_* results in the autocorrelations *ρ_j_* for signal *j* as a list for lags 0 to *n* = ⌊*N/*2⌋, *ρ _j_* = {*ρ*_j0_*, ρ*_j1_, …*, ρ*_jk_, …*, ρ*_jn_}. The autocorrelations’ significance (p-value ≤ 0.01) is assessed by using a list of cutoffs, *ρ _c_* = {*ρ*_c1_*, ρ*_c2_, …*, ρ*_ck_}, determined from a null distribution of autocorrelations for each lag. These null distributions are generated from the calculated autocorrelations of simulated random signals that are created by bootstrapping (re-sampling of the original data with replacement). A signal is categorized in a class corresponding to the lowest lag deemed statistically significant, i.e. in class *Lag l*, where *l* = Min [{*i* : *ρ*_ji_ *ρ*_ci_}], and *i* ∈ 1*,…, k*. The result is a unique classification for each signal into *Lag* classes, which also ensured that any identified autocorrelation at a particular lag cannot possibly arise due to dependence on autocorrelations at lower lags.

The signals that do not show significant autocorrelation at any lag are checked for sudden signal spikes at any time point, and if so classified as spike maxima or minima. For each signal not showing autocorrelation, 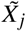 the signal maximum, 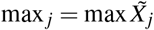, and minimum, 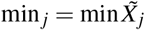, are calculated across all time points. These values are compared against cutoffs {max*_cn_,* min*_cn_*} generated from bootstrap simulated distributions from the data. If for a signal 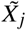 of length *n*, max *_j_ >* max*_cn_* it is classified in the *SpikeMax* class, or otherwise if min *_j_ <* min*_cn_* it is classified in the *SpikeMin* class. Signals for which the signal intensity does not meet cutoff conditions are not reported.

#### Enrichment Analysis

Gene-based over-representation analyses were run using `MathIOmica`^36^ in Mathematica for GO and Reactome database entries. For miRNA enrichment analysis was run using the miRNA Enrichment Analysis and Annotation tool (miEAA) over-representation tool^50^.

Taxa groups were checked for over-representation using `MicrobiomeAnalyst’s`^51^’s web interface. Each subgroup and also each aggregate class were tested on multiple levels. The online MicrobiomeAnalyst database used included the following information, to show analyses at three different levels of mixed-level taxons, species-level and strain-level:

- Mixed-Level Taxon sets included the following taxon sets: 1545 associated with host genetic variations, 239 associated with host-intrinsic factors such as diseases, 118 associated with host-extrinsic factors including diet and lifestyle, 446 associated with environmental factors such as drugs, chemical exposures and 53 associated with microbiome-intrinsic factors such as motility, shape, or spore forming.
- Species-level taxon sets included: 61 associated with host-intrinsic factors including diseases, 92 associated with host-extrinsic factors including diet and lifestyle, 7 associated with environmental factors including drugs and chemical exposures.
- Strain-level taxon sets included: 42 associated with host-intrinsic factors including diseases, 50 associated with microbiome-intrinsic factors such as microbe mobility and shape, and 399 associated with environmental factors including drugs and chemical exposures.

The statistically significant (p *<* 0.05) over-representation results are available with the online data on Zenodo (DOI: 10.5281/zenodo.3358011).

## Network Construction

Weighted expression networks were constructed in which each node represents one molecular species and each edge weight is defined as 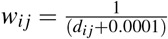, where *d_i j_* is the Euclidean distance between each pair of nodes {*i, j*}, and the offset 0.0001 was added to account for cases where *d_i j_* = 0. Networks were constructed for both the classified daily monitored data and the hourly delta data. To account for missing data in the computation of the Euclidean distance, mean imputation was used. Edge selection for the network construction was determined by filtering on one-tailed quantiles *q*(*N*) based on the *w_i j_* distribution in a given network *k* with *N_k_* nodes:

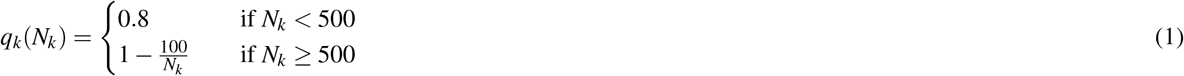

Finally, in the network plots nodes were colored based on the `MathIOmica` classification group to which they belong.

## Acknowledgements

We thank Dr. A. Withrow at the Center for Advanced Microscopy for supporting our TEM; Dr. R. Eby for supporting our NTA. L.R.K.R. was funded through a Bertina Wentworth Endowed Summer Fellowship and the University Enrichment Fellowship at Michigan State University. G.I.M., M.Z., S.D., S.X., C.P., and research reported is supported by Project Number T0412 grant by the Translational Research Institute for Space Health, funded under NASA Cooperative Agreement NNX16AO69A. The content is solely the responsibility of the authors and does not necessarily represent the official views of the supporting funding agencies.

## Author Contributions Statement

G.I.M. conceived of and oversaw the study, carried out project planning, data analysis, data deposition, experimental procedures, and wrote the manuscript. V.V.S. processed experimental samples, prepared protein analysis samples, and prepared small RNA libraries. L.R.K.R. processed samples, analyzed RNA-seq data and assisted in data deposition. G.I.M. and C.P. designed the networks. S.X. coded the network analysis. M.Z. performed data analysis validations. S.D. assisted in network analysis. M.K. performed exosome quantitation and imaging. J.H. carried out experimental protocol design. G.I.M. is the senior author on this work. All authors reviewed the manuscript.

## Competing Financial Interests

G.I.M. has consulted for Colgate-Palmolive North America. C.P. owns equity in Salgomed, Inc. V.V.S., L.R.K.R., M.Z., S.D., S.X, M.K. and J.H. declare the absence of any commercial or financial relationships that could be construed as a potential conflict of interest.

## Notes

https://doi.org/10.5281/zenodo.3358011

https://massive.ucsd.edu/ProteoSAFe/dataset.jsp?task=9c5ded5bda824fd3af1207551901d42a

https://www.ncbi.nlm.nih.gov/geo/query/acc.cgi?acc=GSE108664

https://www.ncbi.nlm.nih.gov/geo/query/acc.cgi?acc=GSE108666

